# Haplotype-resolved *de novo* genome assemblies of four coniferous tree species

**DOI:** 10.1101/2022.11.16.516598

**Authors:** Kenta Shirasawa, Kentaro Mishima, Hideki Hirakawa, Tomonori Hirao, Miyoko Tsubomura, Soichiro Nagano, Taiichi Iki, Sachiko Isobe, Makoto Takahashi

## Abstract

Coniferous trees in gymnosperm are an important source of wood production. Because of their long lifecycle, the breeding programs of coniferous tree are time- and labor-consuming. Genomics could accelerate the selection of superior trees or clones in the breeding programs; however, the genomes of coniferous trees are generally giant in size and exhibit high heterozygosity. Therefore, the generation of long contiguous genome assemblies of coniferous species has been difficult. In this study, we optimized the DNA library preparation protocols and employed high-fidelity (HiFi) long-read sequencing technology to sequence and assemble the genomes of four coniferous tree species, *Larix kaempferi, Chamaecyparis obtusa, Cryptomeria japonica*, and *Cunninghamia lanceolata*. Genome assemblies of the four species totaled 13.5 Gb (*L. kaempferi*), 8.5 Gb (*C. obtusa*), 9.2 Gb (*C. japonica*), and 11.7 Gb (*C. lanceolata*), which covered 99.6% of the estimated genome sizes on average. The contig N50 value, which indicates assembly contiguity, ranged from 1.2 Mb in *C. obtusa* to 16.0 Mb in *L. kaempferi*, and the assembled sequences contained, on average, 89.2% of the single-copy orthologs conserved in embryophytes. Assembled sequences representing alternative haplotypes covered 70.3–95.1% of the genomes, suggesting that the four coniferous tree genomes exhibit high heterozygosity levels. The genome sequence information obtained in this study represents a milestone in tree genetics and genomics, and will facilitate gene discovery, allele mining, phylogenetics, and evolutionary studies in coniferous trees, and accelerate forest tree breeding programs.

## Introduction

Forests cover approximately 31% of the global land area (FAO 2020). In Japan, forest area is as high as more than 60%, of which approximately 40% is accounted for by artificial forests, mainly coniferous trees, which have been planted for applications in the forestry industry (Forestry Agency 2022). *Cryptomeria japonica*, a Cupressaceae family member endemic to Japan, is the most important forestry species in the country, occupying >40% of the artificial forests (Forestry Agency 2022). *Chamaecyparis obtusa*, another member of the Cupressaceae family, is also an important conifer species with superior-quality wood that has been used for the construction of buildings in Japan since the ancient times (Tsumura et al. 2007). *Cunninghamia lanceolata* (Cupressaceae) was introduced from China and Taiwan, and has been recognized as a new woody resource in Japan in recent years because of its fast growth and desirable wood properties (Fujisawa 2017). Additionally, *Larix kaempferi* (Pinaceae) is an important breeding tree species in the northern part of Japan (Kurinobu 2005). Timber tree germplasm and breeding programs have been used to improve wood production and quality. However, the breeding programs of coniferous trees are generally time-consuming because their lifecycle can be as long as or longer than 50 years. Furthermore, the highly heterozygous genomes and outbreeding mating system of coniferous tree species complicate the breeding systems (Burdon and Wilcox 2011).

Since the genomes of coniferous trees are often giant in size (>10 Gb), the analysis of the genome sequence data of these species remains challenging (Neale and Wheeler 2019). Owing to the recent advancements in long-read sequencing technologies, the genomes of gymnosperm species belonging to six families, including Cupressaceae, Cycadaceae, Ginkgoaceae, Pinaceae, Taxaceae, and Welwitschiaceae, have become available (Wan et al. 2022). The giant genomes of gymnosperm species are rich in repetitive sequences such as transposable elements (Ohri 2021). The high-fidelity (HiFi) long-read sequencing technology generates reads spanning the repetitive sequences, resulting in an assembly that covers most of the gene spaces in the genome (Hon et al. 2020).

Coniferous tree genomics using the greatly advanced sequencing technologies promises the acceleration of not only the breeding programs but also the phylogenetic study of coniferous species. In this study, we determined the genome sequences of four coniferous tree species, including *L. kaempferi, C. obtusa, C. japonica*, and *C lanceolata*, using the HiFi long-read sequencing technology. The genome sequence information obtained in this study could serve as a useful resource for breeding coniferous trees, and for understanding the physiology of forest trees and the population genetics and phylogenetics of gymnosperms.

## Materials and methods

### Plant materials and DNA extraction

Four coniferous tree species were used in this study: *Larix kaempferi* (GFE32203) was planted at the Tohoku Regional Breeding Office, Forest Tree Breeding Center, Forestry and Forest Products Research Institute, Forest Research and Management Organization in Iwate, Japan; *Chamaecyparis obtusa* (GFB00119) and *Cryptomeria japonica* (GFA01029) were planted at the Forest Tree Breeding Center, Forestry and Forest Products Research Institute, Forest Research and Management Organization in Ibaraki, Japan; and *Cunninghamia lanceolata* (GFHN00090) was also planted at Forest Tree Breeding Center, Forestry and Forest Products Research Institute, Forest Research and Management Organization in Ibaraki, Japan. The original *C. lanceolata* tree was planted at the Kiyosumi Work Station in the University of Tokyo Chiba Forest in Chiba, Japan.

Genomic DNA was extracted from the young leaves of each tree species using Genome-tips (Qiagen, Hilden, Germany). DNA concentration was measured using the Qubit dsDNA BR assay kit (Thermo Fisher Scientific, Waltham, MA, USA), and DNA fragment length was evaluated by agarose gel electrophoresis with Pippin Pulse (Sage Science, Beverly, MA, USA).

### DNA library preparation and sequencing

DNA libraries were prepared using six protocols (Table 1). Genomic DNA was sheared either by six centrifugations at 1,600 × *g* in the g-Tube (Covaris, Woburn, MA, USA) or in Megaruptor 2 (Deagenode, Liege, Belgium) with the Large Fragment Hydropore mode and mean fragment sizes of 20, 30, or 40 kb. Then, DNA library construction was performed with the SMRTbell Express Template Prep Kit 2.0 (PacBio), according to the manufacturer’s instructions. The obtained DNA libraries were fractionated with BluePippin (Sage Science) to eliminate fragments shorter than 15, 20, or 25 kb in length. The fractionated DNA libraries were sequenced on SMRT cells on the Sequel II and Sequel IIe system (PacBio). HiFi reads were constructed with the CCS pipeline (https://ccs.how).

**Table 1.** HiFi library preparation protocols used in this study

| Protocol | Shearing method | Shearing condition | Library cutoff size |
| --- | --- | --- | --- |
| 1 | g-Tube | Six centrifugations, each at $6,000 \times g$ | 15 kb |
| 2 | g-Tube | Six centrifugations, each at $6,000 \times g$ | 20 kb |
| 3 | g-Tube | Six centrifugations, each at $6,000 \times g$ | 25 kb |
| 4 | Megaruptor 2 | 20 kb | 15 kb |
| 5 | Megaruptor 2 | 30 kb | 15 kb |
| 6 | Megaruptor 2 | 40 kb | 20 kb |

### Genome sequence assembly

Genome sizes of the four tree species were estimated with Genomic Character Estimator (GCE) (Liu et al. 2013), based on the *k*-mer frequency (*k* = 21) calculated with Jellyfish version 2.3.0 (Marçais and Kingsford 2011). The reads were assembled using Hifiasm version 0.16.1 (Cheng et al. 2021) with default parameters. Assembly completeness was evaluated using Benchmarking Universal Single-Copy Orthologs (BUSCO) version 5.2.2, with default parameters (Simão et al. 2015), using the lineage dataset embryophyta_odb10 (eukaryota, 2020-09-10).

## Results

### DNA sequencing

We constructed the DNA libraries of *L. kaempferi, C. obtusa, C. japonica*, and *C. lanceolata* using six library preparation protocols (Protocol1–Protocol6), with slight modification to the sheared DNA fragment sizes and cutoff values to eliminate short DNA in libraries (Table 1). The resultant libraries were sequenced on a total of 38 SMRT cells: 2 cells, Protocol1; 23 cells, Protocol2; 1 cell, Protocol3; 3 cells, Protocol4; 2 cells, Protocol5; and 7 cells, Protocol6.

The number, N50 value, and total length of HiFi reads per cell varied among the protocols (Figure 1). The total length of HiFi reads per cell ranged, on average, from 23.2 Gb in Protocol2 to 35.8 Gb in Protocol6, while the read N50 value of these reads ranged, on average, from 16.3 kb in Protocol4 to 27.8 kb in Protocol6. The read N50 was expectedly long in long-insert libraries prepared using Protocol3 and Protocol6.

**Figure 1.**
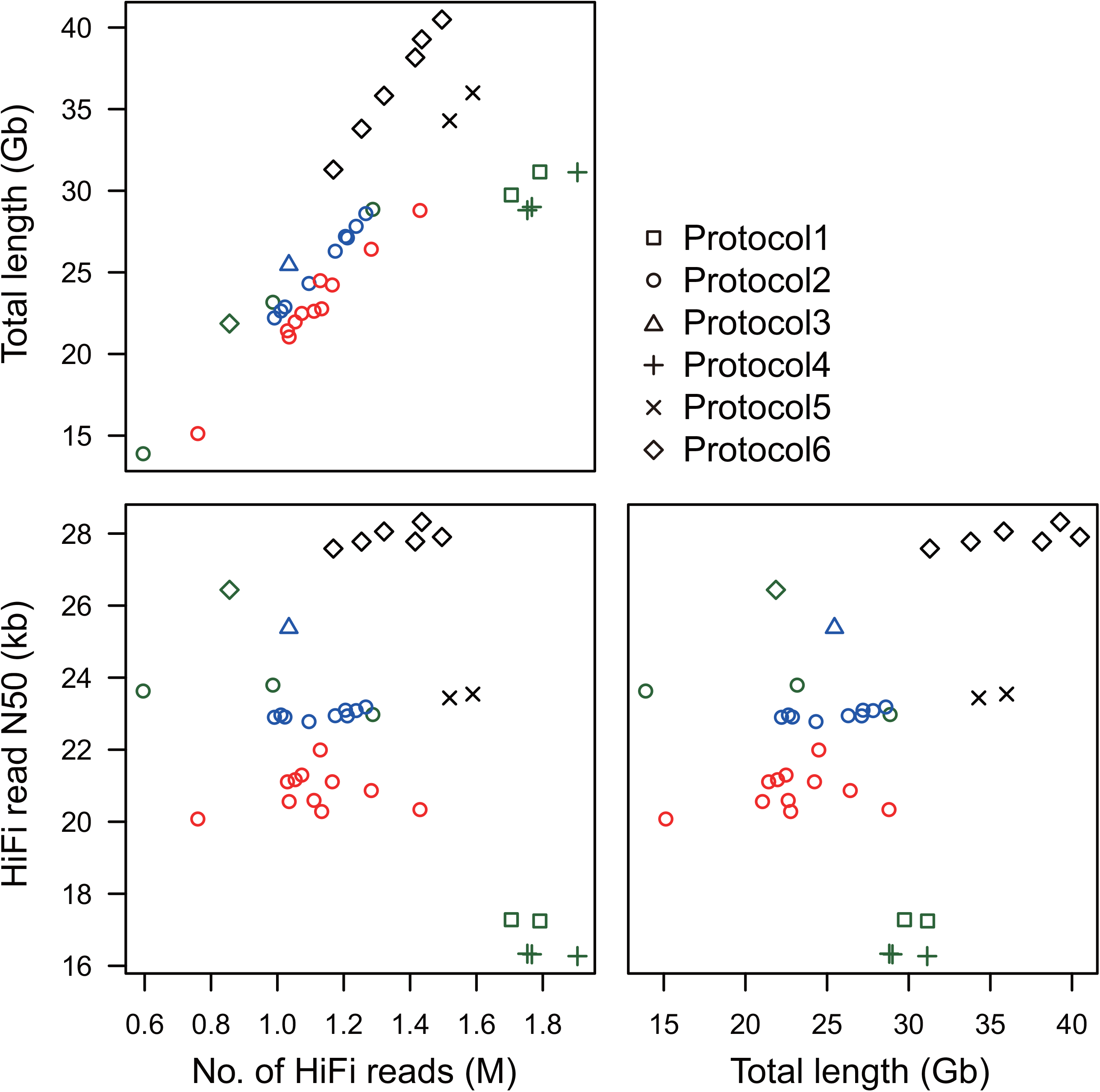
Total length, N50 value, and number of HiFi reads obtained from DNA libraries prepared using six different protocols. The different symbols (square, circle, triangle, plus, cross, and diamond) indicate the different protocols (1, 2, 3, 4, 5, and 6, respectively; detailed in Table 1), and different colors (red, blue, green, and black) indicate the different tree species (*Larix kaempferi, Chamaecyparis obtusa, Cryptomeria japonica*, and *Cunninghamia lanceolata*, respectively).

### Genome assembly of L. kaempferi

A total of 251.4 Gb HiFi reads (median N50 = 20.9 kb) were obtained from 11 SMRT cells (Figure 1). In the *k*-mer distribution analysis, two peaks were detected at *k*-mer multiplicities of 10 and 19 (Figure 2); the former and latter multiplicities corresponded to heterozygous regions of the diploid genome and homozygous regions of haploid genome, respectively. The haploid genome size of *L. kaempferi* was estimated to be 13.2 Gb (Figure 2). Accordingly, the genome coverage of the HiFi reads was calculated as 19× (= 251.4 Gb/13.2 Gb), which was sufficient for *de novo* genome assembly. The reads were assembled with a haplotype-resolved *de novo* assembly method to obtain primary contigs (long continuous stretches of contiguous sequences from one haplotype) and alternate contigs (contiguous sequences including sequence and structural variants from another haplotype). The primary contigs included 4,655 sequences spanning 13.5 Gb, with an N50 value of 16.0 Mb, which corresponded to the estimated size (Table 2). The complete BUSCO score was 89.1% (single-copy BUSCO score = 77.5%, duplicated BUSCO score = 11.6%) (Table 2). On the other hand, the total length of alternate contigs was 9.5 Gb (N50 = 1.7 Mb) (Table 2). The ratio of the length of alternative contigs to that of primary contigs was 70.3% (= 9.5 Gb/13.5 Gb).

**Table 2.**
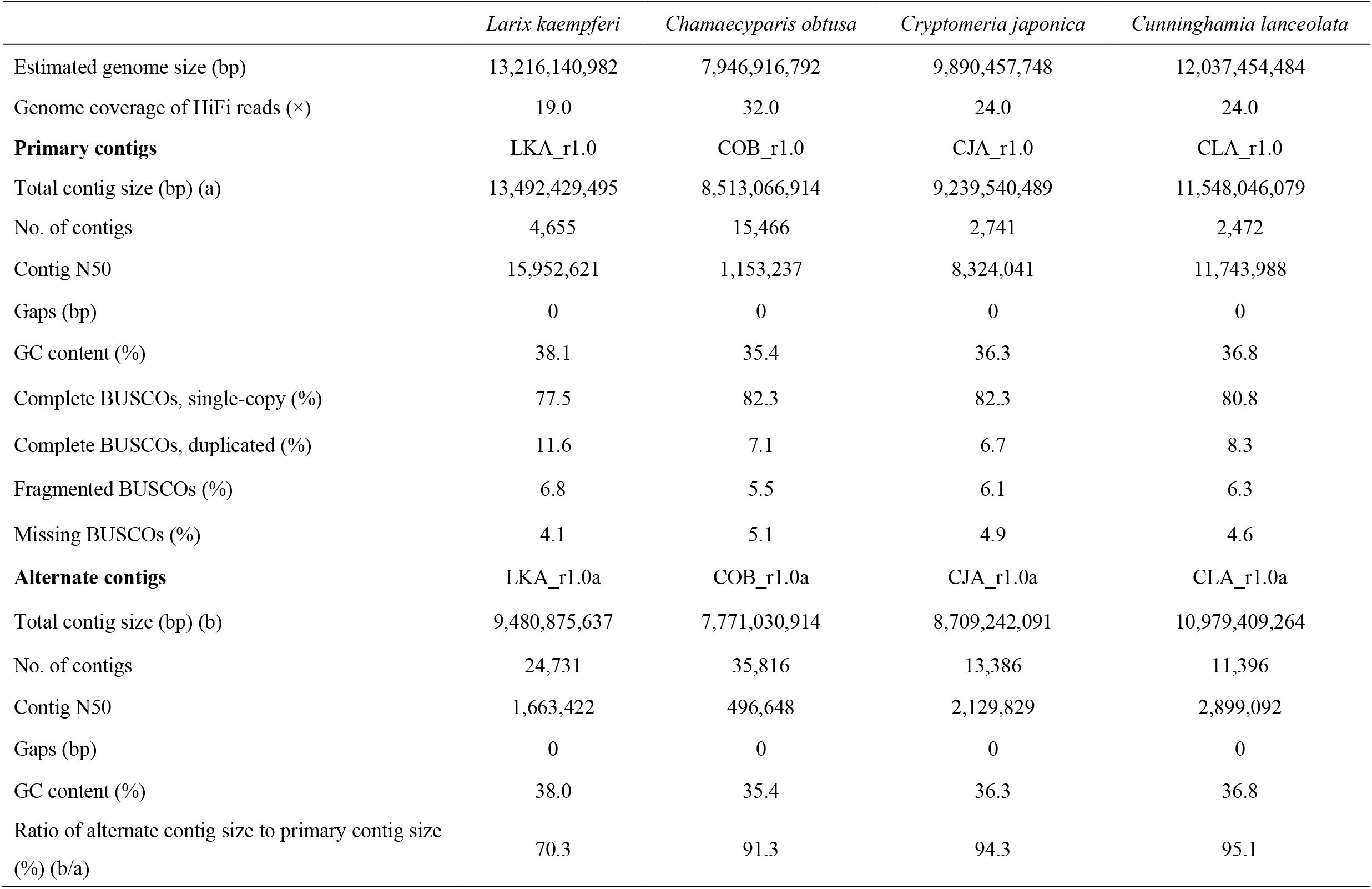
Genome assembly statistics of four timber tree species

|  | *Larix kaempferi* | *Chamaecyparis obtusa* | *Cryptomeria japonica* | *Cunninghamia lanceolata* |
| --- | --- | --- | --- | --- |
| Estimated genome size (bp) | 13,216,140,982 | 7,946,916,792 | 9,890,457,748 | 12,037,454,484 |
| Genome coverage of HiFi reads (×) | 19.0 | 32.0 | 24.0 | 24.0 |
| **Primary contigs** | LKA_r1.0 | COB_r1.0 | CJA_r1.0 | CLA_r1.0 |
| Total contig size (bp) (a) | 13,492,429,495 | 8,513,066,914 | 9,239,540,489 | 11,548,046,079 |
| No. of contigs | 4,655 | 15,466 | 2,741 | 2,472 |
| Contig N50 | 15,952,621 | 1,153,237 | 8,324,041 | 11,743,988 |
| Gaps (bp) | 0 | 0 | 0 | 0 |
| GC content (%) | 38.1 | 35.4 | 36.3 | 36.8 |
| Complete BUSCOs, single-copy (%) | 77.5 | 82.3 | 82.3 | 80.8 |
| Complete BUSCOs, duplicated (%) | 11.6 | 7.1 | 6.7 | 8.3 |
| Fragmented BUSCOs (%) | 6.8 | 5.5 | 6.1 | 6.3 |
| Missing BUSCOs (%) | 4.1 | 5.1 | 4.9 | 4.6 |
| **Alternate contigs** | LKA_r1.0a | COB_r1.0a | CJA_r1.0a | CLA_r1.0a |
| Total contig size (bp) (b) | 9,480,875,637 | 7,771,030,914 | 8,709,242,091 | 10,979,409,264 |
| No. of contigs | 24,731 | 35,816 | 13,386 | 11,396 |
| Contig N50 | 1,663,422 | 496,648 | 2,129,829 | 2,899,092 |
| Gaps (bp) | 0 | 0 | 0 | 0 |
| GC content (%) | 38.0 | 35.4 | 36.3 | 36.8 |
| Ratio of alternate contig size to primary contig size (%) (b/a) | 70.3 | 91.3 | 94.3 | 95.1 |

**Figure 2.**
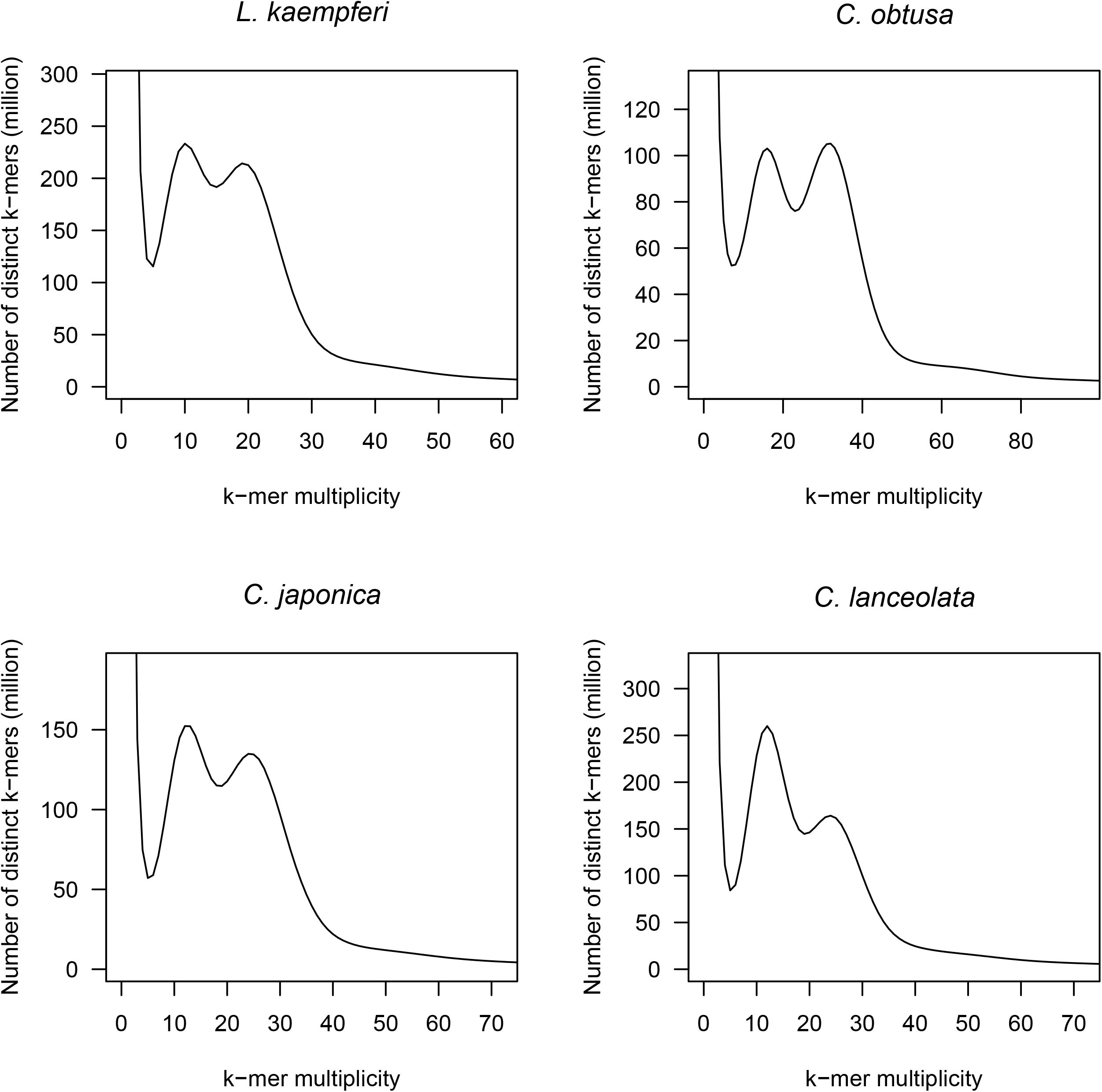
Estimation of the genome sizes of *L. kaempferi, C. obtusa, C. japonica*, and *C. lanceolata* by *k*-mer distribution analysis. The *k*-mer size of 21 was used for the four timber tree species.

### Genome assembly of C. obtusa

A total of 254.5 Gb HiFi reads (median N50 = 23.0 kb) were obtained from ten SMRT cells (Figure 1). Two peaks were detected at *k*-mer multiplicities of 16 (diploid) and 32 (haploid), and the genome size of *C. obtusa* was estimated to be 7.9 Gb (Figure 2). The genome coverage of HiFi reads was 30× (= 254.5 Gb/7.9 Gb). The HiFi reads were assembled into 15,466 primary contigs spanning 8.5 Gb, with an N50 value of 1.2 Mb (Table 2). The total contig length corresponded to the estimated genome size. The complete BUSCO score was 89.4% (single-copy BUSCO = 82.3%, and duplicated BUSCO = 7.1%) (Table 2). Alternate contig length was 7.8 Gb, with an N50 value of 0.5 Mb (Table 2). The ratio of alternate contig length to primary contig length was 91.3% (= 7.8 Gb/8.5 Gb).

### Genome assembly of C. japonica

A total of 237.6 Gb HiFi reads (median N50 = 17.3 kb) were obtained from nine SMRT cells (Figure 1). Two peaks were detected at *k*-mer multiplicities of 12 (diploid) and 24 (haploid), and the genome size of *C. japonica* was estimated to be 9.9 Gb (Figure 2). The genome coverage of HiFi reads was 24× (= 237.6 Gb/9.9 Gb). The HiFi reads were assembled into 2,741 primary contigs spanning 9.2 Gb, with an N50 value of 8.3 Mb (Table 2). The total contig length covered 93.4% of the estimated genome size. The complete BUSCO score was 89.0% (single-copy BUSCO = 82.3%, duplicated BUSCO = 6.7%) (Table 2). The alternate contig length was 8.7 Gb, with an N50 value of 2.1 Mb (Table 2). The ratio of alternate contig length to primary contig length was 94.3% (= 8.7 Gb/9.2 Gb).

### Genome assembly of C. lanceolata

A total of 289.1 Gb HiFi reads (median N50 = 27.8 kb) were obtained from eight SMRT cells (Figure 1). Two peaks were detected at *k*-mer multiplicities of 12 (diploid) and 24 (haploid), and the genome size of *C. lanceolata* was estimated to be 12.0 Gb (Figure 2). The genome coverage of HiFi reads was 24× (= 289.1 Gb/12.0 Gb). The HiFi reads were assembled into 2,472 primary contigs spanning 11.5 Gb, with an N50 value of 11.7 Mb (Table 2). The total contig length covered 95.9% of the estimated genome size. The complete BUSCO score was 89.1% (single-copy BUSCO = 80.8%, duplicated BUSCO = 8.3%) (Table 2).

The alternate contig length was 11.0 Gb, with an N50 value of 2.9 Mb (Table 2). The ratio of alternate contig length to primary contig length was 95.1% (= 8.7 Gb/9.2 Gb).

## Discussion

Here, we report the genome assemblies of four gymnosperm coniferous tree species, *L. kaempferi, C. obtusa, C. japonica*, and *C. lanceolata* (Table 2). Since the genomes of these species were giant in size (∼10 Gb), similar to those of other gymnosperm species, we modified the DNA library preparation protocols to maximize the data production efficiency. Among the six protocols (Table 1), Protocol6 was the most effective with respect to the data yield (median 35.8 Gb) and read N50 length (median 27.8 kb), although the number of reads obtained using Protocol6 was less than that obtained using Protocol4 and Protocol5 (Figure 1). In addition, the genome coverage of HiFi reads differed among the four species. In *C. lanceolata*, HiFi read N50 length was the highest (27.8 kb; Figure 1), and genome coverage was the second highest (24×) among the four species. On the other hand, the HiFi read N50 length of *L. kaempferi* (20.9 kb) was lower than that of *C. lanceolata* (Figure 1), and genome coverage (19×) in *L. kaempferi* was the lowest among the four species. However, unexpectedly, *L. kaempferi* showed the longest sequence contiguity, followed by *C. lanceolata* (Table 2). The heterozygosity level of genomes might affect the assembled sequence contiguity. The ratio of alternate contig length to primary contig length (70.3% in *L. kaempferi* and 95.1% in *C. lanceolata*) might support this assumption. However, the *C. obtusa* assembly was remarkably fragmented, even though its HiFi read N50 (23.0 kb) (Figure 1) and genome coverage (30×) were comparable with those of *C. lanceolata*. This suggests that not only the heterozygosity level but also other genome features, such as the length and/or distribution patterns of repetitive elements, affect the contiguity of the assembled sequence.

In the phylogenetic tree of seed plants, gymnosperms occupy a basal position (Chase et al. 1993) that branches out into the different clades of angiosperms. Therefore, genetic and genomic studies on gymnosperms could shed light on the evolutionary history of plants. However, the genomic investigation of gymnosperms has been lagging behind that of angiosperms because gymnosperms possess large-sized genomes (≥10 Gb) and are relatively less commercially important than angiosperms (Wan et al. 2022). The advent of sequencing technologies has steered this situation in favor of gymnosperm genomics. Long-read sequencing technologies have enabled the sequencing and assembly of giant gymnosperm genomes rich in repetitive sequences, which was not possible with short-read sequencing technologies (Wan et al. 2022). The genome assemblies of four gymnosperm species generated in this study, in addition to those of other species sequenced previously (Wan et al. 2022), will provide new insights into the genome evolution of plants.

The four coniferous species sequenced in this study are important for the forestry. Genome assemblies of these four coniferous tree species could be used as references for the identification of sequence and structural variants in the genomes of divergent cultivars and breeding materials belonging to the same species. Furthermore, in previous genetic and genomic studies, the utilization of a genome sequence as a reference also enabled the identification of genes of interest (Neale and Kremer 2011). The variant and gene information obtained in this study could be used for the development of DNA markers to facilitate genetic studies and breeding programs (Muranty et al. 2014). Genome prediction might be another powerful tool for the selection of elite tree lines from breeding programs (Lebedev et al. 2020; Grattapaglia 2022), which usually require a long time. In addition, gene editing technology could also be used as an effective breeding strategy for shortening the duration of tree breeding programs (Bewg et al. 2018; Goralogia et al. 2021).

The genome sequence information obtained in this study could contribute to breeding programs, gene discovery, and allele mining in coniferous tree species with giant genomes. Since the coverage of genome sequences was as high as ∼90%, according to BUSCO evaluation, the assemblies could be used as reference sequences in transcriptome analysis (Mishima et al. 2022). The protocols presented in this study would contribute to and accelerate the genome sequence analysis of coniferous species with giant genomes. Moreover, chromosome-level assemblies, which would enable phylogenetics and evolutionary studies based on comparative genomics, could also be established through further genomic and genetic analyses in the near future.

## Data availability

Raw sequence reads were deposited in the Sequence Read Archive (SRA) database of the DNA Data Bank of Japan (DDBJ) under the accession numbers DRA014993 (*L. kaempferi*), DRA014992 (*C. obtusa*), DRA014994 (*C. japonica*), and DRA014995 (*C. lanceolata*). The assembled sequences are available at DDBJ (accession numbers: BSBM01000001-BSBM01004655 [*L. kaempferi*]; BSBK01000001-BSBK01015466 [*C. obtusa*]; BSBL01000001-BSBL01002741 [*C. japonica*]; and BSBN01000001-BSBN01002472 [*C. lanceolata*]), BreedingTrees-by-Genes (http://btg.kazusa.or.jp), and Plant GARDEN (https://plantgarden.jp).

## Acknowledgments

We thank the University of Tokyo Chiba Forest for their support and cooperation during the collection of *C. lanceolata* leaf samples. We also thank Y. Kishida, M. Kohara, C. Minami, K. Ozawa, H. Tsuruoka, and A. Watanabe (Kazusa DNA Research Institute) for technical assistance. This study was supported in part by the MAFF commissioned project study on “Development of efficient breeding technique aiming at forestry trees with superior carbon storage capacity” (Grant Number JPJ009841), JSPS KAKENHI (22H05172 and 22H05181), and the Kazusa DNA Research Institute Foundation.

## Competing interests

The authors have no competing interests to declare that are relevant to the content of this article.

## Notes

### Competing Interest Statement

The authors have declared no competing interest.

